# BELIEFS: A Hierarchical Theory of Mind Model based on Strategy Inference

**DOI:** 10.1101/2025.06.25.660690

**Authors:** Giorgio L. Manenti, Sepideh Khoneiveh, Jan Gläscher

## Abstract

Theory of Mind (ToM) refers to the ability to infer another agent’s latent mental states, such as intentions, beliefs, and strategies, to predict their behavior. A core feature of ToM is its recursive structure: individuals reason not only about what others think, but also about what others think about them. Existing computational models typically assume that ToM Level-0 (L0) agents rely on a fixed heuristic (e.g., Win–Stay Lose–Shift, WSLS), an assumption that fails to capture the diversity of non-mentalizing strategies humans actually use. Here we introduce BELIEFS, a probabilistic ToM framework that infers latent L0 strategies directly from behavior using a Hidden Markov Model (HMM) that enables flexible tracking of elemental strategies and dynamic switches between them without relying on a single predefined heuristics. A second HMM tracks the beliefs about the Opponent’s ToM level and dynamic changes therein. We evaluated BELIEFS across four classic dyadic games (Matching Pennies, Prisoner’s Dilemma, Bach or Stravinsky, Stag Hunt) under varying learning rates and volatility of strategy switches. Predictive performance, quantified via cumulative negative log-likelihood (NLL) of the opponent’s choices, was compared against chance and a WSLS-based ToM model, with BELIEFS consistently achieving superior accuracy across conditions. Strategy inference was assessed using trial-wise confusion matrices and Cohen’s κ, revealing robust above-chance classification. Additionally, separability of ToM levels across games indicated that competitive games are particularly informative for distinguishing recursive reasoning from deterministic L0 strategies. The model also successfully tracked the opponent’s recursive reasoning depth (i.e. ToM level) by distinguishing action sequences generated by L0 versus L1 opponents. Parameter-recovery analyses confirmed reliable estimation of core transition parameters. Together, these results show that BELIEFS provides a flexible, computationally grounded account of human ToM, jointly inferring surface-level strategies and recursive reasoning, with applications to modeling adaptive behavior in dynamic interactive environments.

## Introduction

Theory of Mind (ToM) refers to the capacity to infer the unobservable mental states of others ^1–3^, such as their intentions, beliefs, desires, and goals, and to use this information to predict behavior ^4,5^. A central feature of ToM is its recursive nature ^6,7^: individuals can represent not only what others are thinking, but also what others think about themselves (e.g., “I believe that you believe I will do X”). This recursive reasoning is typically organized into hierarchical levels ^8,9^. At Level-0, an agent acts without considering the mental states of others. At higher levels, agents model others as reasoning one level below themselves, forming increasingly complex mental simulations.

ToM is foundational to human social interaction ^10^ and is increasingly important in human-machine contexts, such as collaborative AI ^1^, autonomous agents ^11,12^, and social robotics ^13,14^. At a general level, ToM enables agents to construct mental models of others that support accurate predictions of their future behavior. These models range from non-mentalizing strategies (Level-0) to more sophisticated recursive reasoning (Level-1 and above), where an agent simulates the thought processes of others who, in turn, may be simulating them.

Several computational models have been developed to formalize the hierarchical complexity of mentalizing ^15,16^. These models typically begin with a Level-0 (L0) agent, who knows the rules of the game and selects actions based solely on environmental feedback, without attributing intentionality to the opponent. In this framework, the opponent’s behavior is treated as part of the environment, not as a product of strategic reasoning.

In most ToM models, Level-0 agents are defined using simple and fixed, non-recursive heuristics, such as Win-Stay, Lose-Shift (WSLS), basic reinforcement learning, or frequency-based opponent tracking (e.g., Fictitious Play). WSLS predicts that an agent will repeat a choice following a reward and switch following a loss, which provides a computationally convenient baseline. However, this narrow conception underestimates the variability of human decision-making, even at the most basic levels. For example, in repeated economic games such as the Iterated Prisoner’s Dilemma, reactive strategies such as Tit-for-Tat, which mimic the opponent’s previous move and may be decoupled from the received outcome, can be highly effective without requiring recursive reasoning or Theory of Mind ^17^. These observations highlight a key limitation of many existing models: by relying on static Level-0 assumptions, they fail to capture the richness and adaptability of actual human strategies. Moreover, the choice of Level-0 implementation can influence which higher-order models are favored in model comparison.

Our model addresses this limitation by allowing for flexible L0 strategy inference. Rather than hardcoding a specific heuristic, the model infers the opponent’s L0 strategy from observed behavior and uses this inference as the foundation for building higher levels of reasoning. Specifically, in our framework, a Level-1 (L1) agent who we will refer to with female pronouns (she/her) models the opponent as an L0 agent for whom we will reserve the male pronouns (he/him). She attempts to infer which L0 strategy the opponent is using and then computes the best response to that inferred behavior. Importantly, the best response is calculated on the integration of the beliefs about all possible L0 strategies. At Level-2 (L2), the agent models the opponent as a Level-1 player who, in turn, models the agent as L0. Thus, the L2 agent simulates how her opponent is reasoning about her own behavior and selects a best response to that recursive inference. This iterative structure continues at higher levels, where an agent at Level-(k) simulates an opponent at Level-(k-1), forming a recursive chain of “best responses to best responses.”

This hierarchical organization carries several structural implications for how recursive reasoning unfolds in the model. First, the agent’s mental model of the *opponent is always situated one level below* her own, as each level of reasoning is constructed by computing a best response to the simulated behavior at the lower level. Second, these *recursive simulations are built iteratively*: the agent begins with an assumed Level-0 representation and constructs higher levels step by step by nesting best responses to the previous level’s behavior. Third, and crucial point, the *attribution of Level-0 strategies alternates across levels in a systematic way*. At odd-numbered levels (e.g., Level-1, Level-3), the agent models the hierarchy starting from opponent’s Level-0 strategies, whereas at even-numbered levels (e.g., Level-2, Level-4), the agent considers her own behavior as the base of the levels reasoning (see Figure 1). Finally, because different ToM levels entail different simulated beliefs and best responses, the *model generates distinct predictions of behavior at each level*. These differing predictions are compared to observed choices, enabling inferences about the likely ToM level at which each player is operating.

**Figure 1.**
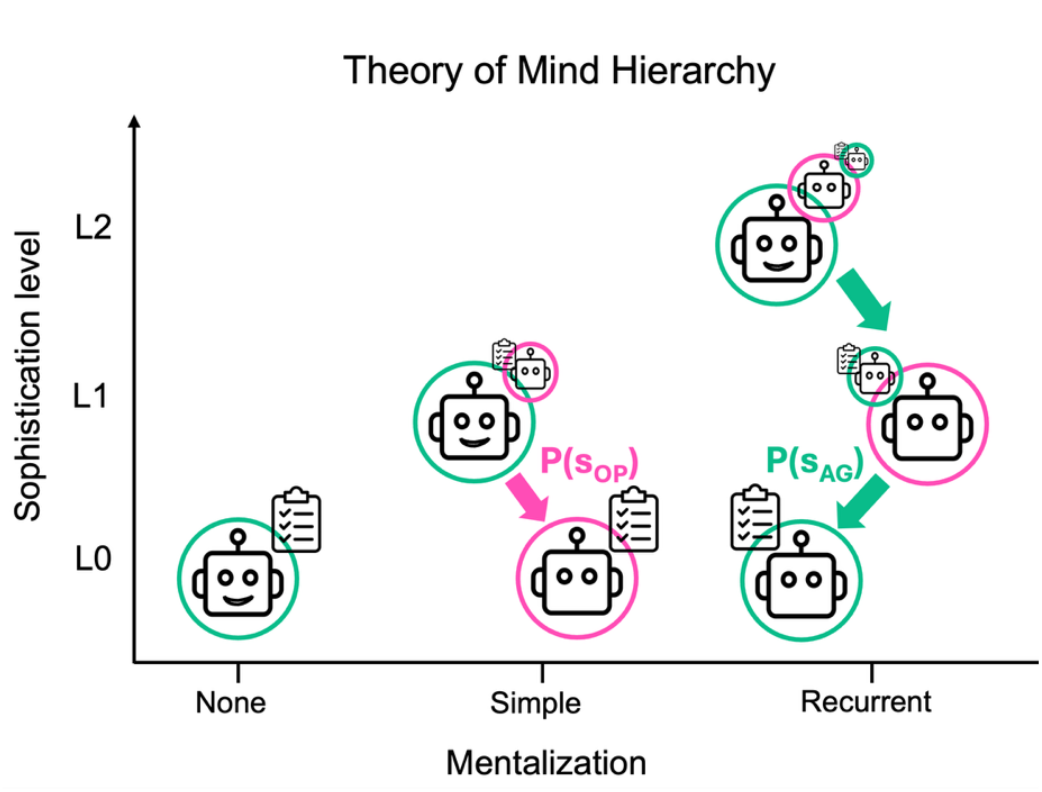
Hierarchy over ToM levels. At Level 0, the agent (AG) follows action strategies without considering the strategies used by the opponent (OP). At Level 1, the agent estimates a probability distribution P(s_OP_) over opponent’s actions, marginalizing over the belief over strategies. At Level 2, the agent simulates the opponent’s best response, assuming the opponent forms a belief about the agent’s strategies P(s_AG_), and then best responds to that. All mentalizing occurs from the agent’s perspective.

Critically, this structure makes the accurate detection of the L0 strategy central to the overall predictive power of the model. All higher-order reasoning depends on this base-level inference. This is the key departure point of our model from earlier recursive ToM frameworks ^15,16^ model Level-0 agents using simple, reward-sensitive strategies, such as Win-Stay/Lose-Shift (WSLS) or reinforcement-learning variants. These implementations provide computationally convenient baselines but do not account for the possibility that human L0 strategies may differ from WSLS or change over time.

Building on this recursive architecture, our framework generalizes traditional ToM models by replacing fixed Level-0 heuristics with a flexible probabilistic inference over multiple candidate strategies. Prior computational ToM models typically assume a single predefined L0 rule, most commonly WSLS or simple stochastic heuristics ^15,16^, or rely on restricted L0 assumptions in agent-based simulations ^18,19^. In contrast, our model maintains a belief distribution over latent L0 strategies and updates these beliefs on every trial using a Hidden Markov Model (HMM). This procedure evaluates how well each strategy’s predicted action explains the opponent’s observed behavior and applies a strategy-transition matrix, parameterized by agent- and opponent-specific stay probabilities, to capture the likelihood of strategy persistence or switching.

These L0 predictions then support a hierarchical best-response process in which each ToM level simulates and responds to the predicted behavior at the level beneath it, following the recursive structure established in earlier ToM modeling frameworks ^15,16^. In parallel, the model tracks the opponent’s depth of reasoning by maintaining and updating a second HMM over ToM levels (Figure 2). After each observed action, the likelihood of that action under competing ToM-level predictions (e.g., but not limited to, L0 vs. L1, Figure 2) is compared, beliefs are updated, and a ToM-level transition matrix regularizes potential level shifts across trials, consistent with recent evidence that mentalizing depth can fluctuate dynamically ^20^.

**Figure 2.**
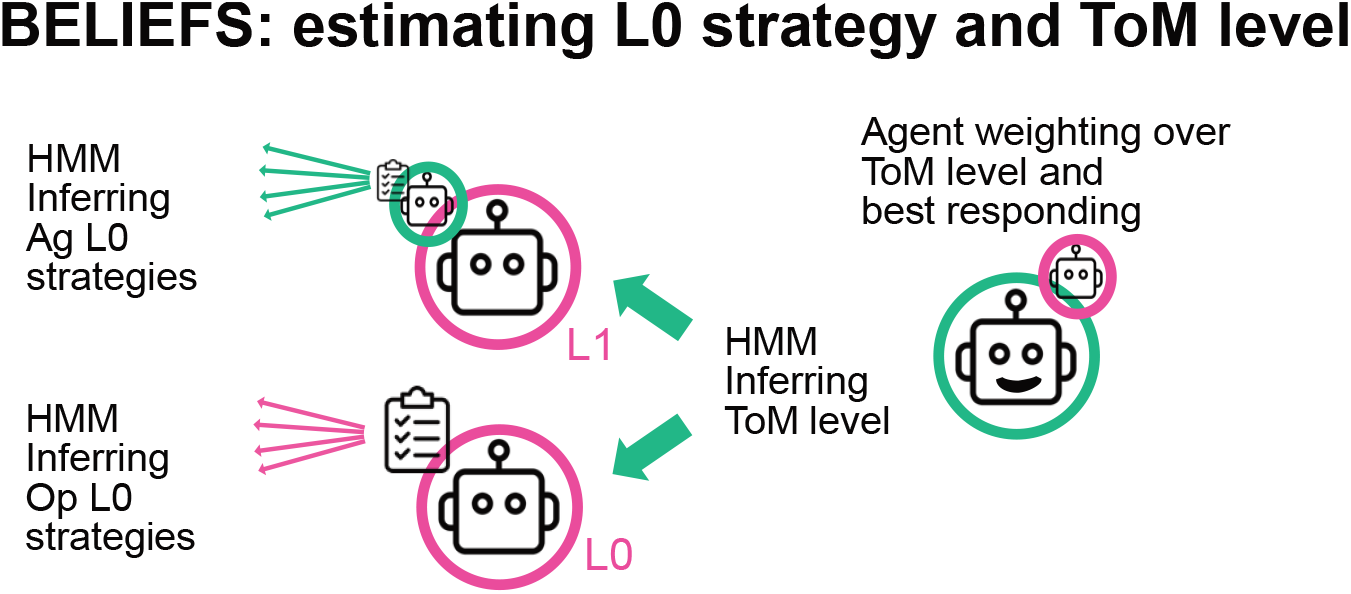
BELIEFS: hierarchical belief inference and action prediction from the perspective of an Agent (green) inferring the Opponent’s ToM level (pink) and Level-0 (L0) strategies. The figure illustrates both the simplified and full recursive structure of the BELIEFS model as implemented by a Level-2 (L2) Agent. To generate these predictions, the model first estimates the L0 strategies of both the Agent and the Opponent (green and pink arrows on the left). This process involves two hidden Markov models (HMMs), each tracking the most probable L0 strategy using a transition matrix parameterized by a stay probability. Both the L0 and L1 Opponent models return action-probability predictions, which are then combined and used to infer the Opponent’s ToM level. This inference is carried out by a third HMM that tracks and updates the probability that the Opponent remains at a given ToM level. Together, the beliefs over ToM level and the corresponding action-probability predictions yield an integrated, level-weighted prediction of the Opponent’s next choice. This prediction is then used by the Agent to compute the best response (see Methods).

This joint inference over surface-level strategies and recursive ToM levels—implemented via continuous Bayesian-style belief updating—constitutes the foundation of our approach, which we term **BELIEFS** (**B**ayesian **E**stimation of **L**evels and **I**ntegration of **E**vidence over **F**lexible **S**trategies). By combining flexible strategy tracking with adaptive inference over reasoning depth, BELIEFS addresses limitations of prior models and provides a unified framework for capturing human strategic cognition in dynamic interactive settings.

## Results

### Predicting L0 Opponent’s action and Tracking L0 strategies

A central aim of Theory of Mind (ToM) models is to predict others’ behavior by inferring their latent mental states, such as beliefs, intentions, and strategies, especially in interactive decision-making contexts. In our study, we evaluate the predictive capabilities of our BELIEFS model by assessing its performance in forecasting an opponent’s next choice under varying levels of environmental volatility. Volatility here refers to the probability that the opponent changes their underlying decision-making strategy (specifically, their Level-0 strategy) on a given trial. We examined a range spanning two theoretically informative extremes: low volatility (5% hazard rate per trial) and high volatility (40% hazard rate per trial).These extremes are chosen in order to approximate plausible human behavioral variability ^21^.

Furthermore, to systematically evaluate model performance, we simulated four classic dyadic games over 40 trials per pair: Matching Pennies, Prisoner’s Dilemma (PD), Bach or Stravinsky (BoS), and Stag Hunt (SH). Each game sequence is simulated across 100 independent dyads.

Predictive performance of the model was assessed using the cumulative negative log-likelihood (NLL) of the Opponent’s choices. Importantly, this analysis focused exclusively on the Level-0 strategy detection component of BELIEFS, without incorporating higher-level recursive ToM inference. The simulated data were generated by truncating the model at the Level-1 Agent, which itself best-responds to a Level-0 Opponent. As a benchmark, we compared our model against both a chance-level baseline and a model that assumes that the opponent follows classical Win-Stay-Lose-Shift at L0. In the WSLS implementation, predicted choices followed a deterministic win/loss rule, passed through a softmax function with a fixed temperature of 1 to ensure consistency with our model’s parametrization.

Across all four game types and under both low and high volatility (hazard rate), our model consistently outperformed the WSLS heuristic in predicting opponent choices (in all games at low volatility; neg. log-likelihood (*BELIEFS-WSLS*) > 24.470, t(49) > 16.718, p <0.0001, Hedges’ g >2.371, FDR corrected; at high volatility; neg. log-likelihood (*BELIEFS-WSLS*) > 21.358, t(49) > 14.572, p <0.0001, Hedges’ g = 2.083, FDR corrected; Figure 3). Importantly, the model always significantly outperformed chance-level prediction (log(0.5)), while this was not the case for WSLS (in all games at low volatility single sided paired test; neg. Log-likelihood (*BELIEFS vs chance*) > 24.119, t(49) > 111.22, p <0.0001, Hedges’ g >15.649, FDR corrected; at high volatility; neg. Log-likelihood (*WSLS vs chance*) < 0.351, t(49) > 0.242, p > 0.959, Hedges’ g <-0.34, FDR corrected; Figure 3).

**Figure 3.**
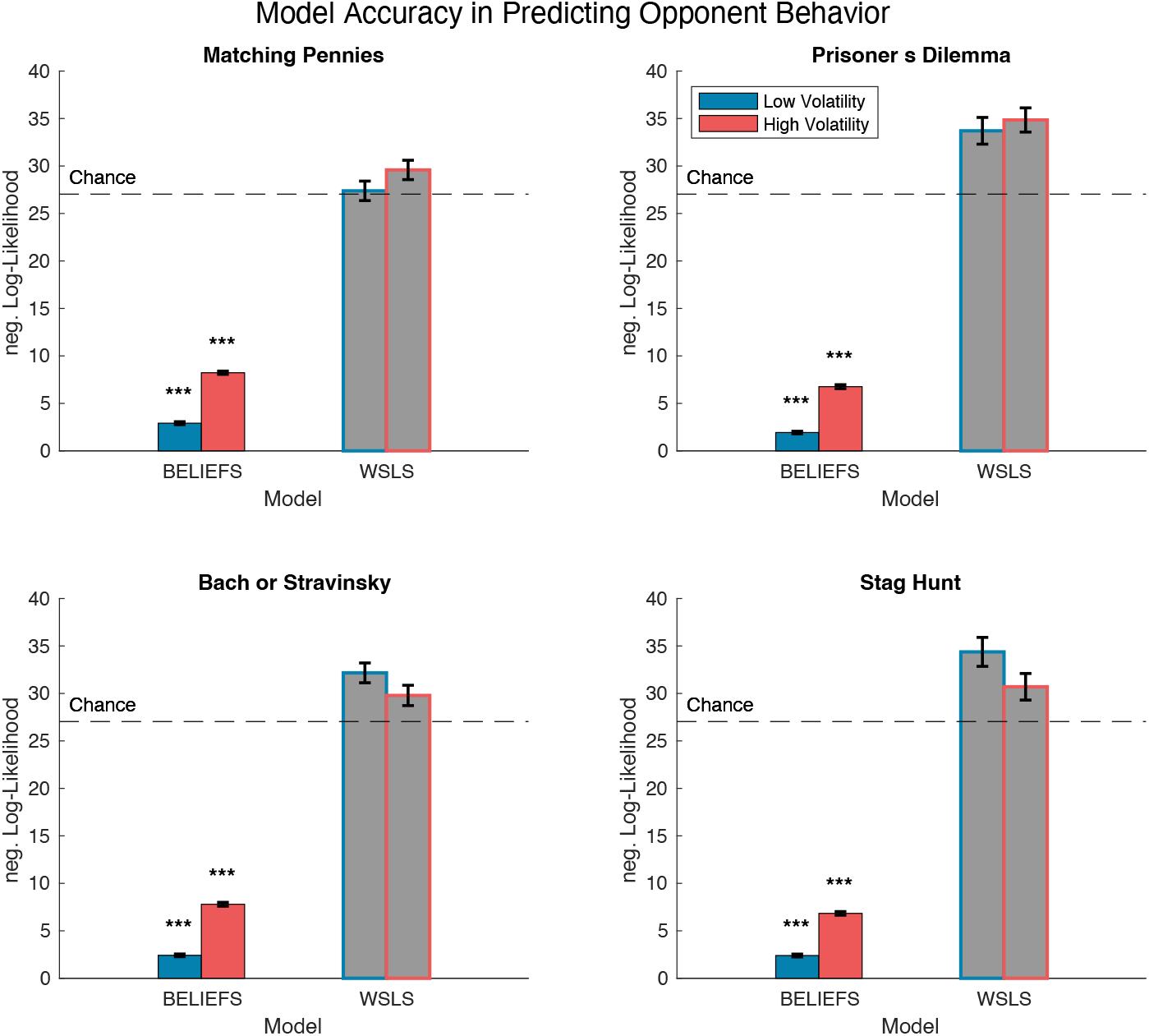
Model Accuracy in Predicting Opponent Behavior. This figure illustrates the predictive performance of the BELIEFS model across four classic dyadic games: Matching Pennies, Prisoner’s Dilemma, Bach or Stravinsky, and Stag Hunt. Each game was simulated over 40 trials for 100 independent dyads. Predictive accuracy is shown for low and high volatility conditions, defined by a median split of the continuous hazard rate, ranging from 5% to 40% probability of Opponent strategy change per trial. Performance is benchmarked against both a classical Win–Stay–Lose–Shift (WSLS) model and chance-level predictions. The WSLS model deterministically predicts the opponent choice based on past action and reward. This predicted action is then converted into a probabilistic prediction using a softmax function with temperature = 1, ensuring direct comparability to the main model. Statistical significance is assessed using paired t-tests against chance performance. Asterisks in the figure indicate conditions where the model significantly outperforms chance (single sided t-test). This analysis highlights the robustness of the model across strategic contexts and volatility levels. In all panels, error bars reflect the SEM. ***p < 0.001, **p < 0.01, and *p < 0.05 (one-tailed t-test, FDR corrected).

Together, these findings demonstrate that the model provides a robust advantage over a WSLS-based ToM heuristic. Full statistics for all comparisons, including means, standard errors, t-values, p-values, and Hedges’ g, are reported in Supplementary Table 1.

Furthermore, beside the significant reliability to predict opponent’s next choice, our model allows to track trial by trial the beliefs about the strategy played by the L0 agent and opponent. In Figure 4, we illustrate the belief dynamics over four L0 strategies, IMITATE, REPEAT, WSLS, and OPPOSE, in the Matching Pennies game under low and high volatility conditions. The plots show how the posterior probability (y-axis) for each strategy evolves over time (x-axis), while the background color denotes the ground truth L0 strategy. These strategy-specific beliefs are used to compute the integrated opponent’ next choice at level-0 *p*(*a*_*OP*_ (*t*)|*S, L*_*OP*_ = 0).

**Figure 4.**
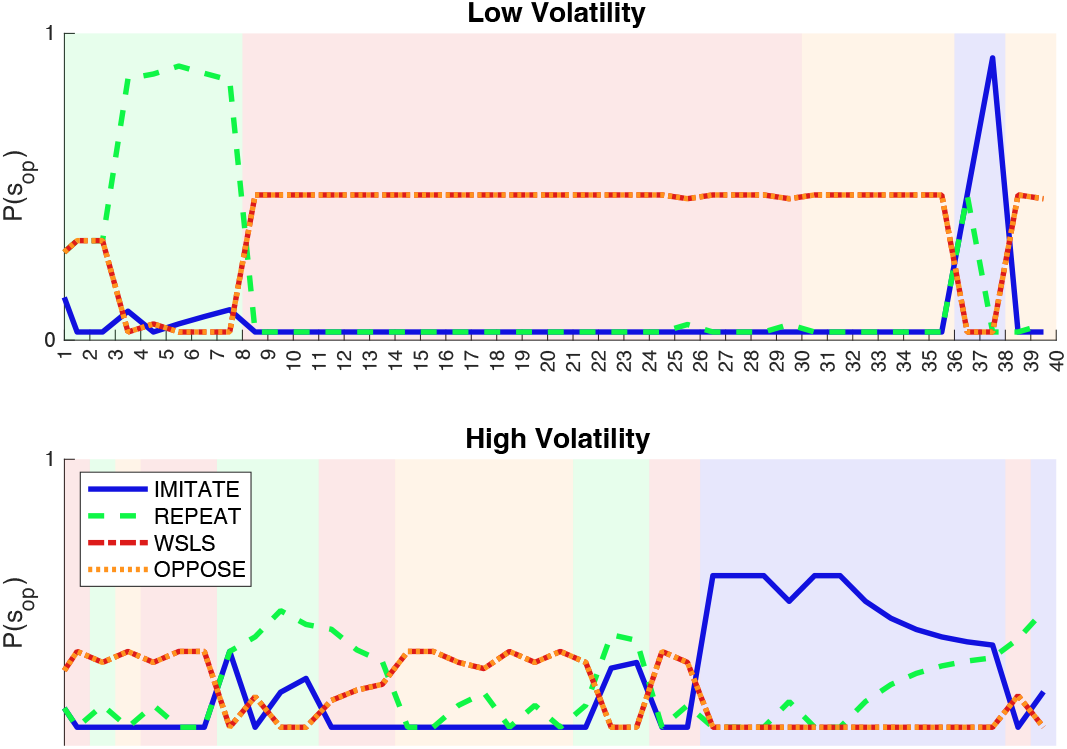
Tracking strategy beliefs. Evolution of the model’s beliefs over four level-zero (L0) strategies, IMITATE, REPEAT, WSLS, and OPPOSE, across 40 trials in the Matching Pennies game. The top panel shows behavior under low volatility, while the bottom subplot shows high volatility. Colored background segments indicate the true underlying opponent strategy on each trial. Strategy belief traces reflect posterior probabilities over time, demonstrating how the model accumulates evidence and updates its beliefs accordingly.

Notably, different latent strategies can produce indistinguishable observable behavior, making pure strategy classification inherently ambiguous in some contexts. For instance, from the perspective of a matcher in MP, WSLS and OPPOSE result always in the same observed actions. This overlap is evident in Figure 4, where the red (WSLS) and orange (OPPOSE) belief traces overlap consistently. However, the model addresses this ambiguity by integrating across beliefs when computing the final opponent action prediction, rather than relying on single-strategy identification. This underscores a fundamental limitation: environments with overlapping behavioral signatures impose structural constraints on how well any model can resolve latent strategies. However, this is not a shortcoming of the model itself. In fact, as shown below in Figure 5, while constrained below a perfect classification the model’s performance approaches fair agreement beyond chance level. To systematically evaluate the model’s ability to infer the correct latent strategy, we computed a trial-by-trial confusion matrix by comparing the model’s predicted strategy (maximum a posteriori) to the true L0 strategy. As a summary statistic, we used Cohen’s Kappa (κ) ^22^, which adjusts for chance agreement (κ = 0 indicates chance performance) and provides a conservative estimate of classification quality (see Methods and Figure 5). Figure 5 displays average κ values across simulated dyads on the four games, stratified by volatility.

**Figure 5.**
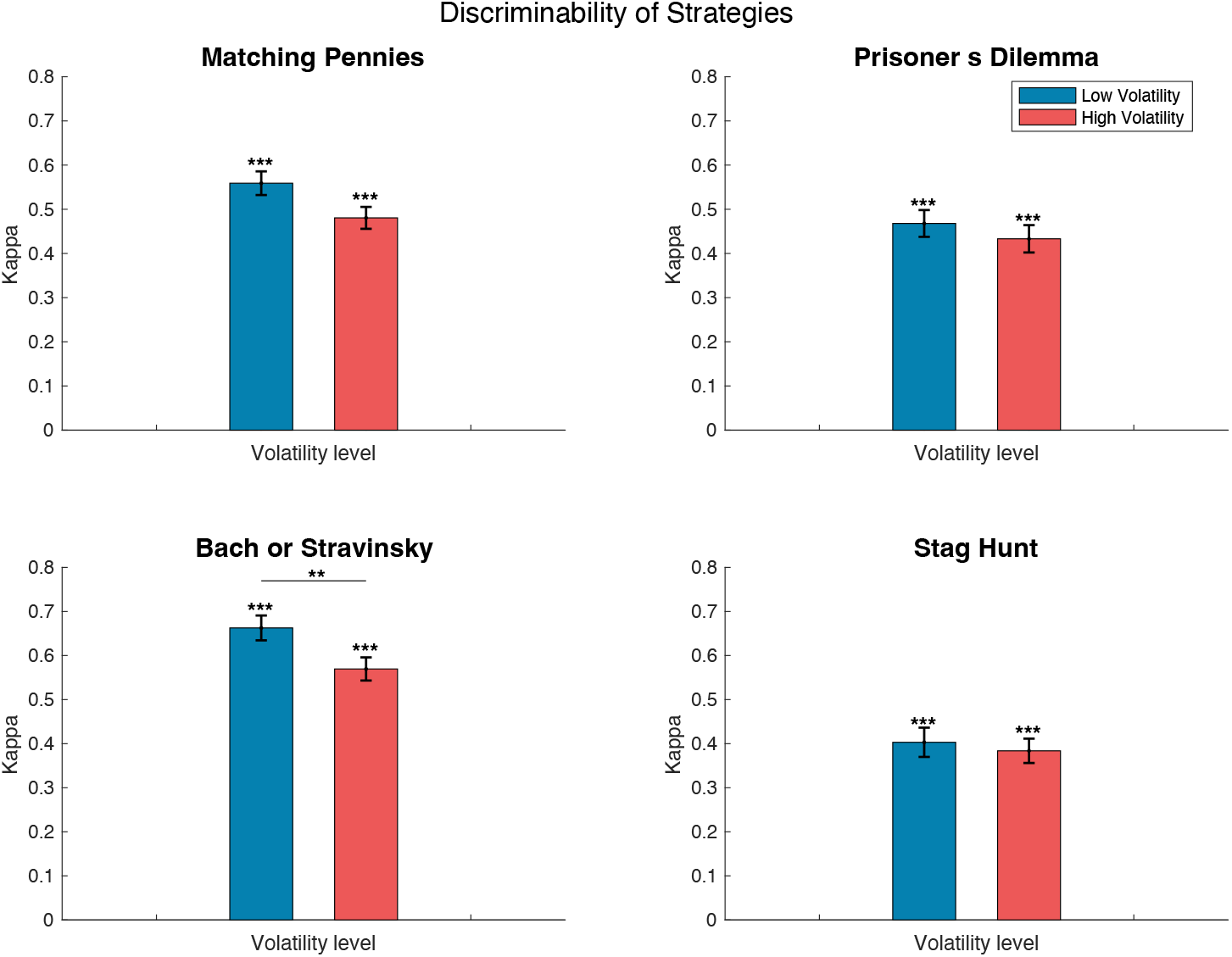
Discriminability of Strategies: Mean Cohen’s Kappa values are shown for four games (MP, PD, BoS, SH), separately for low- and high-volatility conditions (blue vs. red bars). Kappa reflects how accurately the model recovers the true L0 strategy, quantifying agreement between the model-inferred strategy and the ground truth while correcting for chance (κ = 0). Higher values therefore indicate better strategy discriminability. Error bars show ±SEM across participants. Asterisks indicate significance above chance: ***p < 0.001, **p < 0.01, *p < 0.05 (one-tailed t-test, FDR corrected).

Under low volatility, the model reliably tracked the underlying L0 strategy across all games. In all four games, κ values were significantly above zero with lowest bound in SH (mean κ = 0.40, t(41) = 9.060, p < 0.0001, Hedges’ g = 1.37). Similarly, under high volatility, the model’s ability to infer the correct strategy was reduced but still in fair agreement (mean κ = 0.38, t(49) = 16.282, p < 0.0001, Hedges’ g = 2.267). Importantly, all positive Kappa values were significantly above chance, and values exceeding 0.38 are typically interpreted as reflecting at least fair to moderate agreement beyond chance in classification tasks ^23^. Under low volatility, the model’s performance consistently approached or exceeded this threshold, indicating meaningful to substantial tracking of latent strategies. However, while these results show difference over games even between volatility levels, they suggest the BELIEFS can be generalized over games and volatility levels without losing reliability.

### Distinguishing opponent ToM levels and disentangling agent level

To quantify how accurately BELIEFS infers the opponent’s recursive depth of reasoning, the simulation framework described earlier was extended. For each game, 40 action sequences were generated for 100 agent–opponent pairs. As in previous analyses, volatility was manipulated by allowing the opponent’s Level-0 (L0) strategy to switch with either a low (∼10%) or high (∼40%) trial-wise probability. Critically, the opponent’s ToM level (L0 vs. L1) was allowed to switch independently at the same rate, enabling an assessment of whether the model can track both the opponent’s surface-level strategy and their latent ToM level. BELIEFS infers ToM level by comparing the likelihood of the observed opponent action under the L0 and L1 generative models, tracking level beliefs through a Hidden Markov Model that incorporates both Bayesian belief updating and accounting for assumed switches in ToM levels. The resulting trial-wise ToM level beliefs can be appreciated in Figure 6.

**Figure 6.**
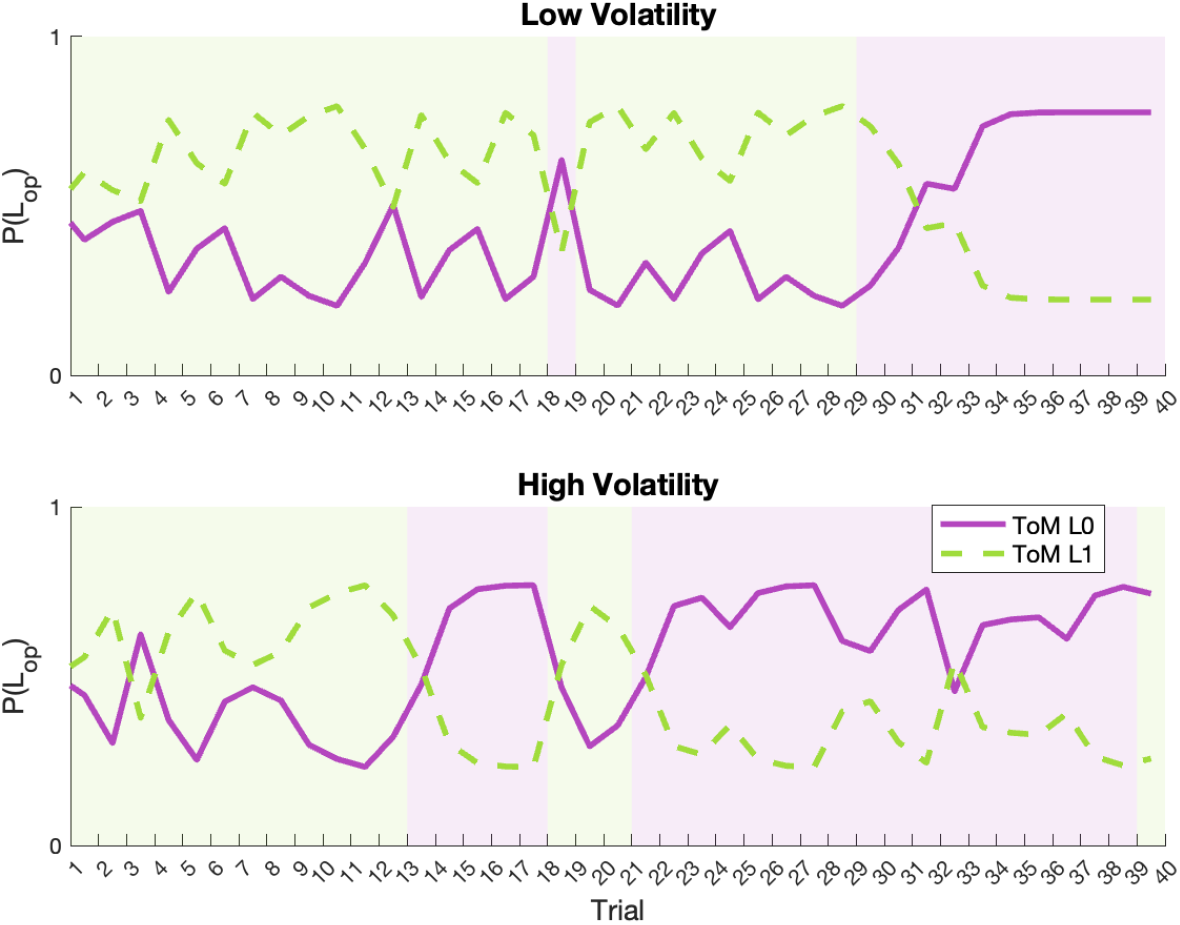
Tracking ToM level beliefs. Evolution of the model’s beliefs over opponent’s ToM level *P(L*_*op*_), either no ToM, level 0 or ToM1, across 40 trials in the Matching Pennies game. The top panel shows behavior under low volatility (10% probability of switching strategy per trial), while the bottom subplot shows high volatility (40% switching probability). Colored background segments indicate the true underlying opponent ToM level on each trial. ToM level belief traces reflect posterior probabilities over time, demonstrating how the model accumulates evidence and updates its beliefs accordingly.

To systematically evaluate the model’s ability to infer the correct latent ToM level, we computed a trial-by-trial confusion matrix by comparing the model’s predicted strategy (maximum a posteriori) to the true L0 strategy. Across all games and conditions, the model reliably assigned lower NLL to actions belonging the true ToM level than to the incorrect one. In all four games, κ values were significantly above zero with lowest bound in BoS (mean κ = 0.30, t(49) = 9.659, p < 0.0001, Hedges’ g = 1.34). Similarly, under high volatility, the model’s ability to infer the correct strategy was reduced but still fair agreement (mean κ = 0.34, t(49) = 10.964, p < 0.0001, Hedges’ g = 1.53) (Figure 7).

**Figure 7.**
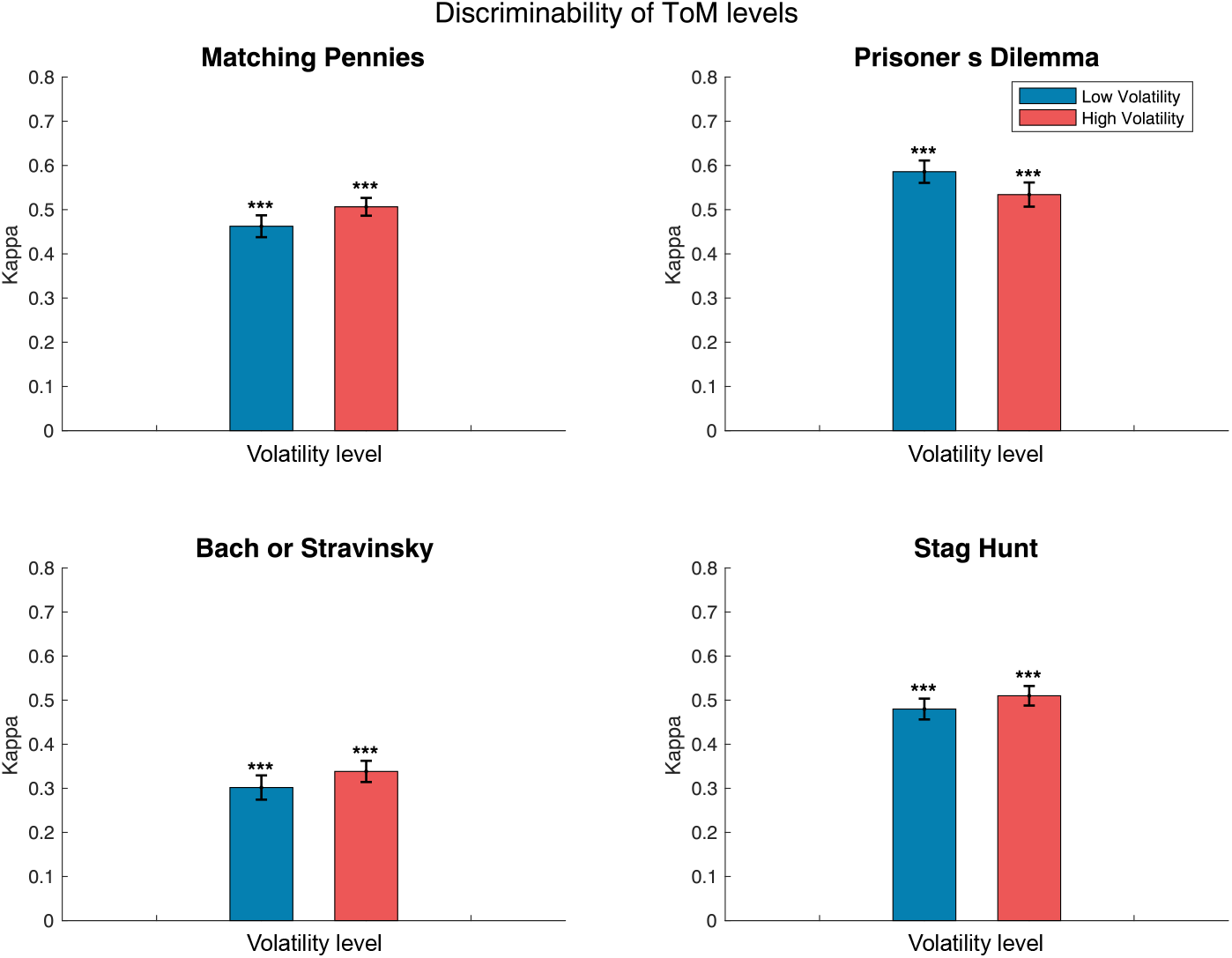
Discriminability of opponent’s TOM level: Average Cohen’s Kappa values are shown across four games (MP, PD, BoS, SH), stratified by learning rate (x-axis) and volatility condition (bar color: blue = low, orange = high). Kappa quantifies agreement between the model’s inferred L0 strategy and the ground truth, corrected for chance (κ = 0). Error bars represent ±SEM across simulations. Asterisks denote significance against chance level: ***p < 0.001, **p < 0.01, *p < 0.05 (one-tailed t-test, FDR corrected).

Next, we evaluated the model’s ability to disentangle the agent’s own ToM level, a key feature for fitting behavioral data. Specifically, we measured the dissimilarity between model-generated action probabilities for agents at ToM level-0 *p*(*a*_*AG*_ (*t*)|*S, L*_*AG*_ = 0), level-1 *p*(*a*_*AG*_ (*t*)|*S, L*_*AG*_ = 1), and level-2 *p*(*a*_*AG*_ (*t*)|*S, L*_*AG*_ = 2), using Fisher-transformed correlations across trials. Greater dissimilarity reflects more distinctive behavioral profiles across levels, supporting reliable inference of the agent’s reasoning depth.

As shown in Figure 8 dissimilarity scores were significantly greater than zero for most pairwise comparisons (p < 0.001, one-tailed t-tests against 0, FDR-corrected), particularly in competitive games. However, in cooperative tasks (BoS and SH), agents at level 0 and level 2 often produced highly correlated behaviors, resulting in lower, but still significant, dissimilarity. This suggests that cooperative contexts may reduce the distinguishability of ToM levels, as different agents may converge on similar strategies despite differing reasoning processes.

**Figure 8.**
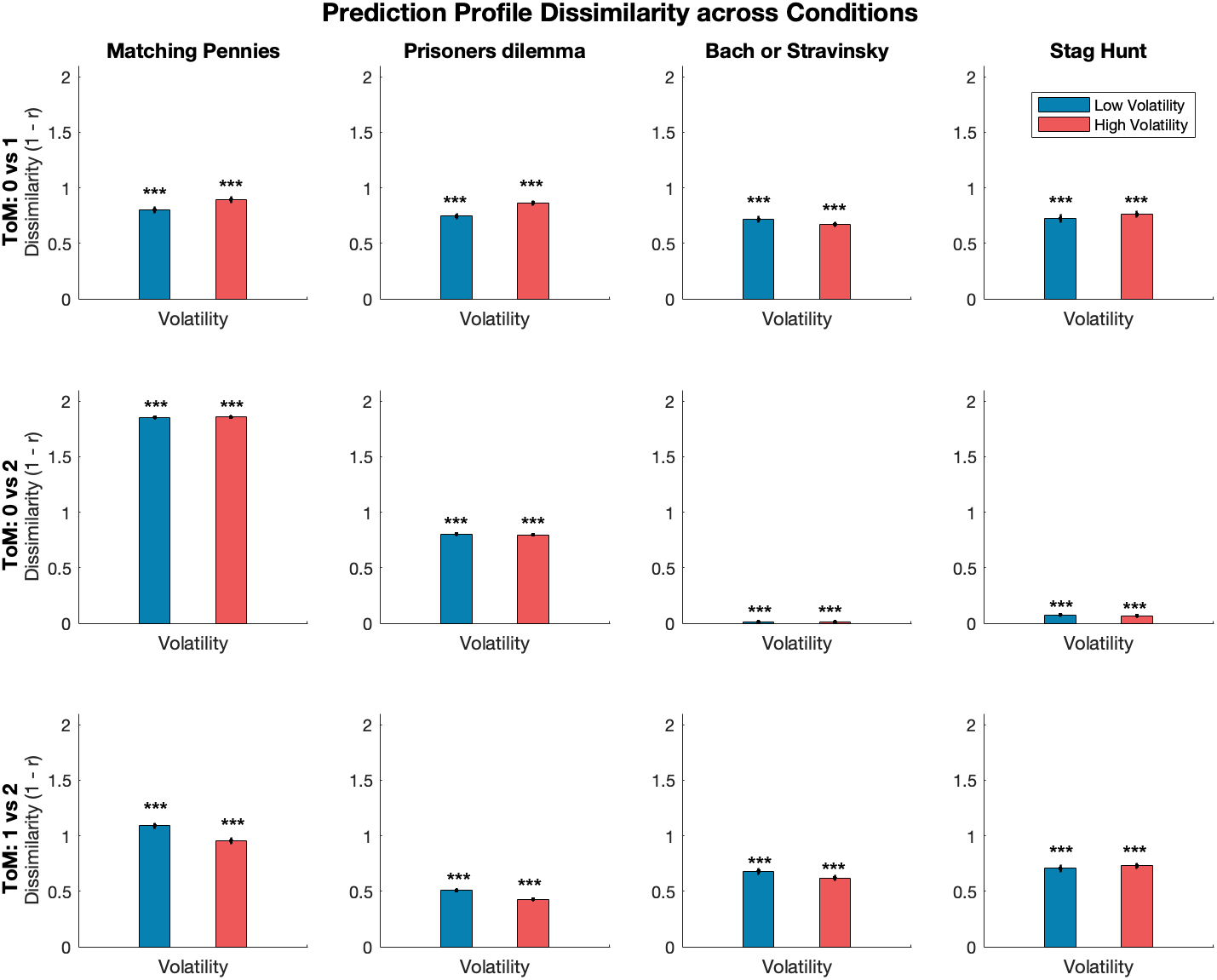
Separability of Theory of Mind levels across games. To evaluate the BELIEFS model’s ability to distinguish between different Theory of Mind (ToM) levels, we computed the dissimilarity between trial-wise action probability distributions for agents at ToM levels 0, 1, and 2 (*p*(*a*_*AG*_ (*t*)|*S, L*_*AG*_ = 0), l *p*(*a*_*AG*_ (*t*)|*S, L*_*AG*_ = 1), *p*(*a*_*AG*_ (*t*)|*S, L*_*AG*_ = 2),). This measure captures the theoretical separability of reasoning levels based on observed behavior across four matrix games. In competitive games such as Matching Pennies (MP), dissimilarity scores were consistently high, indicating that actions at different ToM levels are strongly distinguishable. In particular, ToM0 and ToM2 agents often exhibited opposite (anticorrelated) behavior, producing dissimilarity values approaching the maximum, reflecting the ability to detect recursive reasoning. By contrast, in coordination games like Prisoner’s Dilemma (PD), Bach or Stravinsky (BoS), and Stag Hunt (SH), dissimilarities, especially between ToM0 and ToM2, were markedly lower. This convergence arises because, in these games, higher-level reasoning often produces actions similar to those generated by deterministic L0 strategies, making disentangling ToM levels more challenging. In BoS, for instance, ToM0 and ToM2 agents produced nearly indistinguishable behavior. Despite these differences, all pairwise dissimilarities were significantly greater than zero (p < 0.001, FDR corrected), indicating that some level of behavioral differentiation remains detectable. Error bars represent ±SEM across participants. These results suggest that competitive games with opposing incentives are particularly suitable for observing and distinguishing recursive ToM behavior, whereas coordination games may obscure such distinctions.

**Figure 9.**
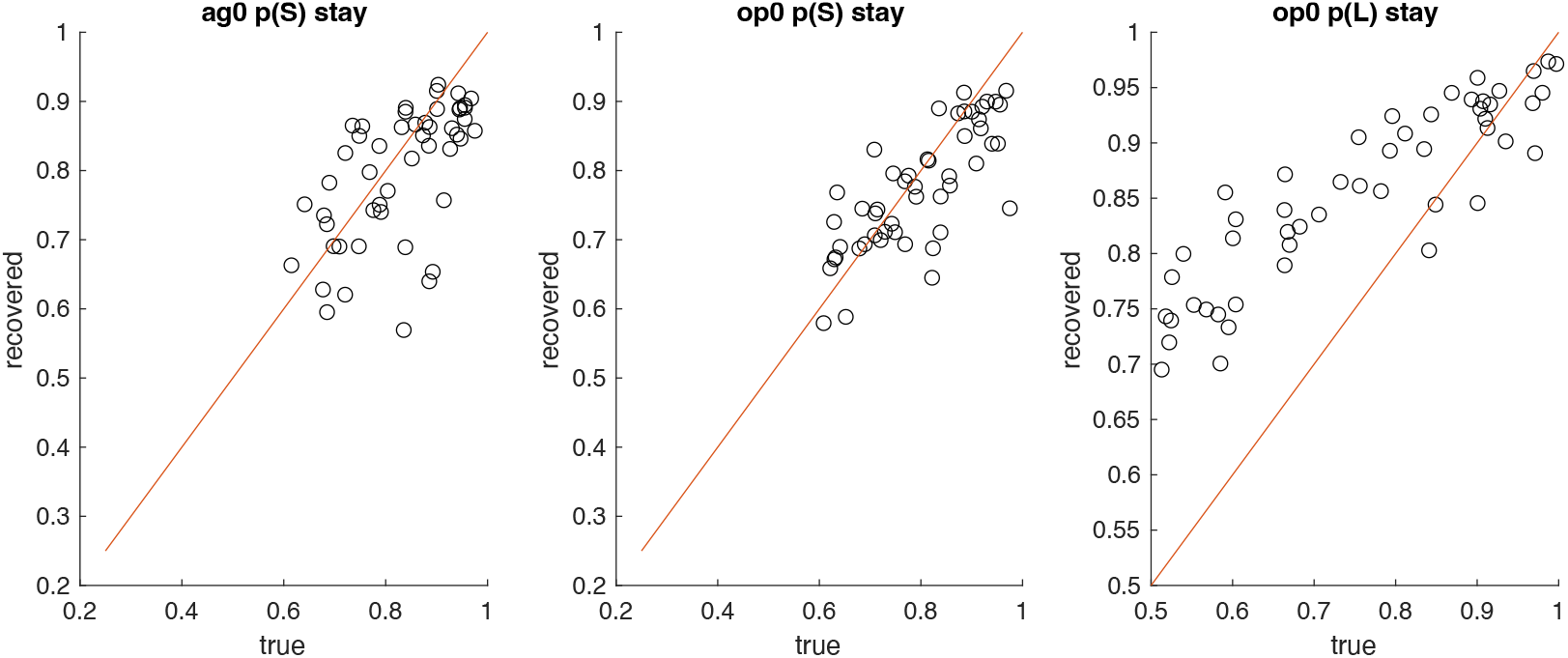
Parameter Recovery: True (x-axis) versus recovered (y-axis) parameter values for the BELIEFS model across 500 simulated datasets are shown. Panels display the Agent’s strategy-stay probability (ag_stay, left), the Opponent’s strategy-stay probability (op_stay, middle), and the Opponent’s ToM-level stay probability (op_pl, right). Diagonal lines indicate perfect recovery (y = x), and points above or below the diagonal reflect over- or underestimation. While ag_stay and op_stay were recovered with high accuracy, op_pl exhibited systematic overestimation for values below ∼0.7, within the limits corresponding to the possible uniform transition matrices (0.25 for strategy-stay; 0.5 for ToM-level stay).

Together, these results show that the model can accurately infer both the opponent’s and the agent’s level of recursive reasoning. This dual capacity to track latent opponent ToM levels and differentiate between internally generated reasoning levels is a key strength of the model, particularly under stable, competitive conditions.

### Parameter recoverability

To assess the validity of the BELIEFS model, we conducted a parameter-recovery analysis. 50 combinations of transition-matrix parameters, namely strategy-stay probabilities for the Agent and Opponent (ag_stay, op_stay) and the ToM-level stay parameter (op_pl), were randomly sampled from uniform distributions (stay parameters: 0.6–1.0; op_pl: 0.5–1.0). For each combination, 10 synthetic action sequences were generated, yielding a total of 500 datasets. The model was fitted to each dataset using the CBM toolbox to maximize the likelihood of the agent- and opponent-integration terms (agInt, opInt; see Methods), with parameter fitting constrained to ranges corresponding to the possible uniform transition matrices (e.g., 0.25–1.0 for strategy-stay, 0.5–1.0 for op_pl). Recovered parameters showed good correspondence with the ground-truth values for ag_stay and op_stay, with moderate-to-strong correlations between true and recovered values (ag_stay: r = .60, p < .001; op_stay: r = .77, p < .001) and regression slopes not significantly different from 1. In contrast, recovery of the ToM-level stay parameter op_pl was less accurate. Although true and recovered values remained highly correlated (r = .89, p < .001), values below approximately 0.7 were systematically overestimated, resulting in a positive bias (mean bias = +0.12). Overall, these results indicate that the model parameters are generally recoverable, with minor overestimation for low values of the ToM-level stay parameter.

Importantly, across these simulated datasets, the model’s probability estimate of the opponent’s ToM level (op_pl) correlated reliably with the true ToM level (mean r = .48, 95% CI [.45, .50], p < .0001; Fisher z-averaged). This indicates that the model systematically tracks whether the opponent is L0 or L1.

## Discussion

Theory of Mind (ToM) is not a unitary process ^24^, but rather a family of socio-cognitive mechanisms that support the attribution of first-order beliefs and values to others, integrating these representations into one’s own decision-making. In this work, we present a model that advances current approaches by relaxing a key constraint: the fixed Level-0 (L0) strategy assumption, which is often instantiated using the Win-Stay-Lose-Shift (WSLS) heuristic ^15,16^. Instead, our framework allows for probabilistic inference over a broader set of opponent strategies, enhancing both the accuracy of ToM-level estimation and the predictive validity of the model. By moving beyond rigid heuristics such as WSLS and adopting a distributional approach to strategy and ToM-level inference, the model better captures how humans might reason about others. Prior models frequently rely on rational agent assumptions that presume optimality or fixed rules, assumptions often violated in real-world behavior ^25,26^. In contrast, our framework embraces behavioral variability and supports bounded rationality by enabling agents to infer, weight, and adapt to a range of possible opponent strategies and reasoning depths.

A further limitation is that the current evaluation relies primarily on simulated data rather than human behavior. Although BELIEFS incorporates dynamic updating of strategy and ToM-level beliefs, empirical validation is needed to determine how well these mechanisms generalize to real social decision-making. Our separability analysis also showed that the discriminability of ToM levels is task-dependent. In cooperative games such as Bach-or -Stravinsky and Stag Hunt, Level-0 and Level-2 agents often exhibit highly correlated behavior, reducing, though not eliminating, dissimilarity between levels. This suggests that cooperative structures may mask differences in recursive reasoning because successful coordination can emerge without explicit mentalizing. If human behavior reflects similar mechanisms, competitive tasks may offer more diagnostic leverage for identifying ToM-level differences, unless differences between cooperation and competition constitute the central research question.

A central strength of the proposed architecture is its scalability. The framework can, in principle, be applied to any matrix game structure, including multi-agent and dynamic scenarios, without requiring modifications to the inference mechanism. While our model offers a promising framework for scalable and cognitively plausible ToM reasoning, it has thus far been evaluated only on simulated data. It will also be important to examine how social reasoning interacts with affective or moral components of cognition currently outside the scope of our model.

Notably, our framework is structured to support inference over both the agent’s and the opponent’s ToM levels, enabling the tracking of recursive beliefs, not only about others’ strategies but also about their reasoning depth. This opens a promising direction for future work, particularly in addressing long-standing debates about whether human mentalizing operates via discrete levels or more continuous gradients of belief. Moreover, such probabilistic outputs could, in principle, be aligned with neural data (e.g., functional imaging) to study how social representations evolve over time during decision-making. In future work, we aim to more directly evaluate the model’s inference over ToM levels by computing trial-by-trial posterior probabilities of each level given the opponent’s action.

Beyond theoretical implications, flexible ToM tracking has substantial applications. In human-AI interaction, systems that can infer human goals and social reasoning depth can respond more adaptively in collaborative and mixed-initiative contexts, such as assistive robotics or digital companions ^1^. Similarly, in education, such models could underpin adaptive learning systems that infer student misunderstandings not only from observable errors but also from inferred reasoning strategies and beliefs ^27,28^. In both domains, dynamic and probabilistic ToM modeling bridges computational inference with the variability and nuance of human cognition.

In summary, we introduce a novel, flexible model of Theory of Mind that accumulates evidence over multiple strategies and integrates hierarchical beliefs to compute best-response action probabilities. By relaxing rigid prior assumptions and dynamically tracking social reasoning depth, the model provides a scalable and cognitively informed foundation for future research in decision science, neuroscience, and human-AI interaction.

## Material and Methods

### Software implementation and availability

All codes and simulation data are available for download on GIN. All data simulation and analyses were carried out in Matlab 2024a (The Mathworks) Statistics for p-value correction was done with fdr_bh toolbox for matlab ^29^ following the ^30^ method.

### L0 Strategies definition and set choice

Level-0 (L0) strategies in the model are defined as simple heuristics that generate deterministic predictions based on recent contextual information; namely, the most recent actions and outcomes observed in the game. These strategies operate without any Theory of Mind (ToM) reasoning and treat the opponent’s behavior as an environmental regularity rather than the product of intentional decision-making. In the present study, we consider a fixed set of four candidate L0 strategies commonly discussed in the literature on repeated matrix games (e.g., Matching Pennies, Prisoner’s Dilemma; ^31–33^). However, it is important to note that the model is not limited to this set, additional strategies can be included or excluded depending on the context of application. The four strategies used are defined as follows:

#### IMITATE

This strategy predicts that the opponent will repeat the agent’s last choice (*c*_*OP*_ (*t* + 1) = *c*_*AG*_ (*t*)). In the literature on the Prisoner’s Dilemma game, this strategy is known as Tit-for-Tat.

#### REPEAT

This strategy predicts that the opponent is simply repeating his last choice (*c*_*OP*_ (*t* + 1) = *c*_*OP*_ (*t*)). This strategy partially overlaps with the winning part of WSLS.

#### OPPOSE

This strategy is the opposite of IMITATE and predicts that the opponent will oppose the agent and deliberately not choose the agent’s last choice (*c*_*OP*_ (*t* + 1) ≠ *c*_*AG*_ (*t*))

#### WSLS (Win-Stay, Lose-Shift)

This strategy predicts that the opponent will repeat his current choice following a win in the current trial (*c*_*OP*_ (*t* + 1) = *c*_*OP*_ (*t*) | *win*), whereas he will shift to the other choice option following a loss in the current trial (*c*_*OP*_ (*t* + 1) ≠ *c*_*OP*_ (*t*) | *loss*).

Note that while these strategies only evaluate the current choices of Agent and Opponent, it is also possible to include strategies defined on the sequence of choices in recent trials.

### L0 belief integration

The model for an L1 agent cycles over a finite set of strategies *s* ∈ *S* and evaluates whether the current choice of the L0 opponent matches each strategy and accumulates for it. We consider that the L0 Opponent is following a certain strategy *s* ∈ *S* for a number of trials and switch to another strategy with an unknown hazard rate *h*. These strategies serve as the hidden states in a Hidden Markov Model (HMM) and they will emit choice predictions in the form of binary action probabilities. Currently, all strategies evaluate past choices and outcomes to derive a discrete choice prediction for the current trial, also known as a (binary) strategy-specific belief over choices. These take the form *p*(*a*_*OP*_(*t*)|*s*) = [1 0] if *c*_*OP*_ = 1 or *p*(*a*_*OP*_(*t*)|*s*) = [0 1] if *c*_*OP*_ = 2.^1^ For instance, the *IMITATE* strategy predicts that *c*_*OP*_(*t* + 1) = *c*_*AG*_(*t*), so if *c*_*AG*_(*t*) = 1, then *p*(*a*_*OP*_(*t*)|*s* = *IMITATE*) = [1 0].

We can construct these L0 Opponent action probabilities for all possible strategies in the set and collect them in a prediction matrix *p*(*a*_*OP*_(*t*)|*S*) with the number of strategies and actions as the dimensions. Although the strategies make discrete action predictions, the strategies themselves are unobservable latent variables that must be inferred from observable choices. In most cases, when the action predictions of two strategies overlap, the L1 agent cannot know exactly which strategy the L0 agent is following. In this situation, the optimal L1 agent will weigh and integrate these predictions with beliefs that reflect how often each strategy has made matching action predictions in the past.

Thus, the beliefs *p*(*s*_*OP*_) (probability weight for each strategy of the L0 Opponent) play a central role for the L1 agent for calculating the integrated action predictions for the L0 Opponent:

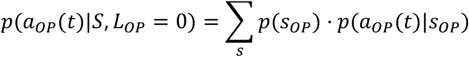

Here, we explicitly condition the integrated action prediction on the level of the opponent, because this differentiation will become important, when constructing the ToM hierarchy below. The agent can then best-respond to these integrated action predictions (see below).

### Updating of strategy beliefs using Bayesian belief updating

Having derived the matrix of action probabilities *p*(*a*_*OP*_(*t*)|*S*), the observed choice of the Opponent *c*_*OP*_(*t*) can be used to calculate the likelihood of the observed choice *p*(*a*_*OP*_ = *c*_*OP*_(*t*)|*S*). To obtain the posterior (updated) strategy beliefs *p*(*S*|*c*_*OP*_(*t*)), can then be obtained via Bayesian updating by multiplying the likelihood with the prior beliefs and normalizing to ensure that the posterior remains a valid probability distribution:

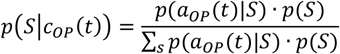

Within the framework of a Hidden Markov Model, the posterior beliefs must also account for potential transitions to other strategies. This is achieved by incorporating a transition matrix *T*_*s*_ which specifies the probability of moving from strategy *s* to *s*′ on the next trial:

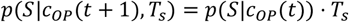

The transition matrix *T*_*s*_ can be constructed in several ways. The simplest variant, which we use here, estimates the stay probability *p*_*stay*_ from the data during model fitting and sets this probability along the diagonal *T*_*s*_ indicating the common probability to maintain the same strategy in the next trial. All off diagonal elements will be assigned the identical switch probability (or hazard rate) *p*_*switch*_ = (1 − *p*_*stay*_)/*n*_*s*_ − 1, where *n*_*s*_ is the number of strategies in *S*. This adjusted posterior strategy belief *p*(*S*|*c*_*OP*_(*t*), *T*_*s*_) will then serves as the prior for the next trial.

### Best responding to L0 Opponent predicted actions

Consistent with previous computational theory-of-mind (ToM) models ^15,16,19^, the player being modeled (here referred to as the Agent) is defined as one level above the player she is modeling (i.e., the Opponent). In the BELIEFS model, the action at the higher level is assumed to be a best response to the predicted actions of the lower-level player ^20,34^. Specifically, the model computes the expected value of each choice option and then applies a softmax function to derive the action probabilities for the Agent.

The expected value of a choice is defined in the standard way as the product of reward magnitude and reward probability:

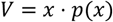

In the context of matrix games with joint actions (*c*_*AG*_, *c*_*OP*_) and outcomes (*U*_*AG*_, *U*_*OP*_) for both players, the Agent receives only the outcome associated with her chosen action *c*_*AG*_, weighted by the probability that the Opponent selects a specific action *c*_*OP*_. Here, *a*_*OP*_ refers to the set of all possible actions available to the Opponent, while *c*_*OP*_ denotes the concrete action actually selected on a given trial. Consequently, the expected value for the Level-1 Agent is

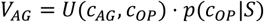

and the corresponding action probabilities are computed as

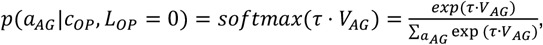

where τ denotes the inverse temperature parameter controlling choice stochasticity.

In summary, to best-respond to the action predicted at the lower ToM level, the Agent needs to have an accurate estimate of those action probabilities at the lower level *p*(*a*_*OP*_|*S*). The goal of the BELIEFS model, as with other computational ToM models, is to estimate these probabilities as accurately as possible.

### Considering different ToM levels for the Opponent

When the Opponent is simulated as a Level-1 player, the Agent correspondingly operates at Level 2, reasoning about the Opponent’s Level-1 best response to the integrated strategy predictions over the Agent’s Level-0 behavior that the Opponent is assumed to track. A Level-1 Opponent mirrors the calculations of a Level-1 Agent, but from the opposite perspective. Specifically, the Level-1 Opponent evaluates the Agent’s choices as a Level-0 player and estimates the strategy beliefs of the Level-0 Agent. It is important to note that strategy predictions for a Level-0 Agent generally differ from those of a Level-0 Opponent due to differences in underlying choices. Consequently, beliefs over the strategies of a Level-0 Agent are maintained separately from those of a Level-0 Opponent.

The Level-1 Opponent constructs the strategy prediction matrix for the Level-0 Agent, *p*(*a*_*AG*_(*t*)|*S*), in the same manner described previously, and integrates these predictions with the strategy beliefs of the Level-0 Agent, *p*(*s*_*AG*_), according to

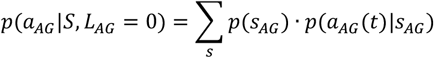

The Level-1 Opponent then computes a best response to the integrated action probabilities of the Level-0 Agent:

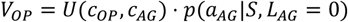

Where *U*(*c*_*OP*_, *c*_*AG*_) refers to the Opponent’s joined payoff matrix, because the Opponent’s choice *c*_*OP*_ is used as the first argument. These values are then converted into action probabilities using a softmax function:

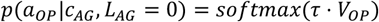

The Level-2 Agent subsequently uses these calculations to best respond to the predicted actions of the Level-1 Opponent:

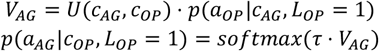

### Updating of beliefs about the Opponent’s ToM level

The BELIEFS model also maintains and updates a belief over the theory-of-mind (ToM) level of the Opponent using the same Hidden Markov Model (HMM) inference mechanism as applied to Level-0 strategies. Predicted action probabilities for the Opponent at different ToM levels can be conceptualized as the emission probabilities of the hidden states, which correspond to the ToM levels. These probabilities are derived either from the integrated strategy predictions of a Level-0 Opponent, *p*(*a*_*OP*_(*t*)|*S, L*_*OP*_ = 0), or from the best-response calculations of a Level-1 Opponent to the Level-0 Agent, *p*(*a*_*AG*_|*c*_*AG*_, *L*_*AG*_ = 0).

Weighting the Opponent’s action predictions with the beliefs over ToM levels, *p*(*L*_*OP*_), produces level-adjusted action predictions, organized in a matrix with ToM levels and actions as its dimensions:

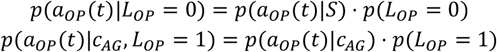

These level-adjusted action predictions are used in two ways. First, marginalizing over ToM levels by summation yields the overall predicted action probabilities for the Opponent, *p*(*a*_*OP*_(*t*)), which the Level-2 Agent can best respond to:

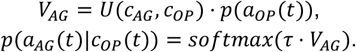

These predictions differ from those computed previously, as they incorporate beliefs over ToM levels. Second, the level-adjusted action predictions are used to update ToM beliefs. Conditioning on the actual Opponent choice *c*_*OP*_(*t*) and normalizing over levels produces the posterior ToM level beliefs:

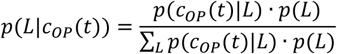

The action probabilities *p*(*a*_*OP*_(*t*)|*L*) have already been multiplied with the prior ToM level beliefs, completing the Bayesian update in two steps.

Finally, the beliefs over ToM levels are adjusted for potential switches in the Opponent’s level, analogous to strategy shifts at Level-0. A transition matrix over ToM levels, *T*_*L*_, is defined, with the diagonal representing the probability of staying at the same level, *p*(*L*′ = *L*), estimated from the data during model fitting. All non-diagonal elements are assigned the uniform switch probability (1 − *p*(*L*′ = *L*))/(*nL* − 1), where *nL* is the number of ToM levels considered:

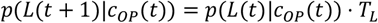

These updated ToM level beliefs then serve as the prior for the next trial, completing the HMM-like updating of ToM beliefs.

### Prediction of opponent choice analysis

To evaluate how well the BELIEFS model predicts Opponent choices under different internal learning dynamics and environmental volatility, we generated simulated datasets from a Level-1 Agent interacting with a Level-0 Opponent. The Opponent’s strategy switches were governed by two volatility regimes, implemented as random switching probabilities with a low-volatility mean of 0.08 and a high-volatility mean of 0.31. These values reflect an underlying continuous distribution of switch rates, but for analysis we grouped simulations into low versus high volatility conditions using the same generating distributions. For each simulation, the model’s trial-wise beliefs were used to compute the probability assigned to the Opponent’s actual choice, yielding a log-likelihood for every trial (excluding the first). Summing these values produced a subject-wise negative log-likelihood (NLL), which served as the primary accuracy metric. To benchmark performance, we compared BELIEFS to a Win–Stay Lose–Shift (WSLS) heuristic. WSLS deterministically repeats the previous action after a win and switches after a loss; to ensure comparability, we transformed WSLS predictions into probabilistic form using a softmax policy with temperature = 1 (equivalent to an effective scaling factor of 0.269 on WSLS choice tendencies). For both the BELIEFS model and WSLS, we computed trial-wise and participant-wise NLLs and conducted one-sided t-tests against chance-level log-likelihoods. To directly compare models, paired t-tests were used to test whether BELIEFS achieved higher predictive accuracy than WSLS. Effect sizes are reported as Hedges’ g, corrected for sample size, and all p-values were adjusted using false-discovery-rate (FDR) correction (Benjamini & Hochberg, 1995). All reported p-values were corrected for multiple comparisons using the FDR procedure (Benjamini & Hochberg, 1995). This procedure allowed us to quantify how model performance varies as a function of both internal learning parameters and environmental volatility, and to determine whether the BELIEFS model reliably outperforms a baseline heuristic.

### Strategies confusion matrix analysis

To evaluate how well the model discriminates among latent strategies, we computed Cohen’s kappa ^22^ between the model-predicted and true strategies across trials. This measure corrects for chance agreement and provides a normalized metric of classification performance, with κ = 1 indicating perfect prediction and κ = 0 corresponding to chance-level agreement. Formally, Cohen’s kappa is defined as:

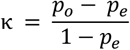

where *p*_*o*_ is the observed proportion of agreement and *p*_*e*_ is the expected proportion of agreement by chance. For each trial, the model outputs a posterior probability distribution over strategies, denoted as P(s), where s indexes the strategy space. To generate discrete predictions, we selected the strategy with the maximum posterior. In cases of ties (i.e., multiple strategies with equal maximum probability), we resolved them by randomly sampling one of the tied strategies. To account for the stochasticity introduced by random tie-breaking, we repeated this procedure 100 times and computed a confusion matrix and kappa coefficient for each sample. The final model-based kappa was obtained by averaging across bootstraps.

To determine whether performance exceeded chance, we conducted one-sample t-tests comparing subject-level κ values against zero (chance level), correcting p-values for multiple comparisons using the FDR procedure. Confusion matrices were computed only for dyads in which more than one strategy was expressed over time; sequences containing a single, unchanging true strategy (e.g., 3.5% of MP game sequences) were excluded from the analysis.

For each subject, we constructed a normalized confusion matrix by comparing predicted and true strategies across trials. Rows were normalized to sum to one, representing conditional probabilities of predicting each strategy given the true label. We then averaged these confusion matrices across subjects and bootstrap samples to compute the overall confusion matrix. Kappa was computed from this averaged confusion matrix using standard formulae for multiclass classification.

### Parameter recovery

Fifty different combinations of generating parameters governing the transition matrix (strategy stay parameters: *p*_*switch*_ for ag_stay, op_stay; and theory-of-mind level stay parameter: op_pl) were randomly sampled from uniform distributions ranging from 0.6 to 1.0 for the stay parameters and from 0.5 to 1.0 for op_pl. For each of the 50 parameter combinations, 10 sequences were generated. Using the CBM toolbox, the model was fitted to maximize the likelihood of integrated agent and opponent actions. The fitted transition matrices ranged between a uniform transition matrix (all transition probabilities equal) and a sharp diagonal matrix (probability of 1.0 to stay in the same state). The average recovered parameters were compared to the true generating parameters. Likewise, the average model likelihood based on the generating parameters (computed from the action probabilities of the generating sequence) was compared to the recovered likelihood. The softmax temperature was fixed at 1.0 during generation, and similar results were obtained when varying the softmax temperature.

### Disentangling agent levels

We quantified the dissimilarity between model predictions across ToM levels by computing the pairwise dissimilarity of the action-wise log-likelihoods predicted by the model under different ToM levels (0, 1, and 2). For each simulated agent, we constructed a 3×3 Fisher-z-transformed correlation matrix between the predicted action log-likelihoods for each pair of ToM levels. These correlation matrices were averaged across trials and then converted into dissimilarity values as:

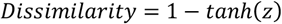

where z is the Fisher-z-transformed correlation between two ToM levels. We extracted the dissimilarity for each pair (0 vs 1, 0 vs 2, 1 vs 2) and analyzed how it varied across model parameters (learning rates) and task conditions (volatility).

### Parameter recovery

To validate the ability of the BELIEFS model to recover generating parameters, synthetic datasets were generated using the model itself. For each parameter, values were sampled from uniform distributions: stay parameters for the Agent and Opponent (ag_stay, op_stay) ranged from 0.6 to 1.0, and the ToM-level stay parameter for the Opponent (op_pl) ranged from 0.5 to 1.0. Fifty unique combinations of these generating parameters were sampled. For each combination, 10 action sequences were generated, yielding a total of 500 datasets. Each dataset was subsequently fitted using the CBM toolbox. Fitting was performed by minimizing the composite negative log-likelihood of the integrated action probabilities for both the Agent and the Opponent (agInt and opInt; Figure 10C), given the corresponding generated actions. Recovered parameters were then compared to the true generating values to assess the accuracy of recovery. Correlation coefficients, regression slopes, and mean bias were computed to quantify the agreement between true and recovered parameters.

**Figure 10.**
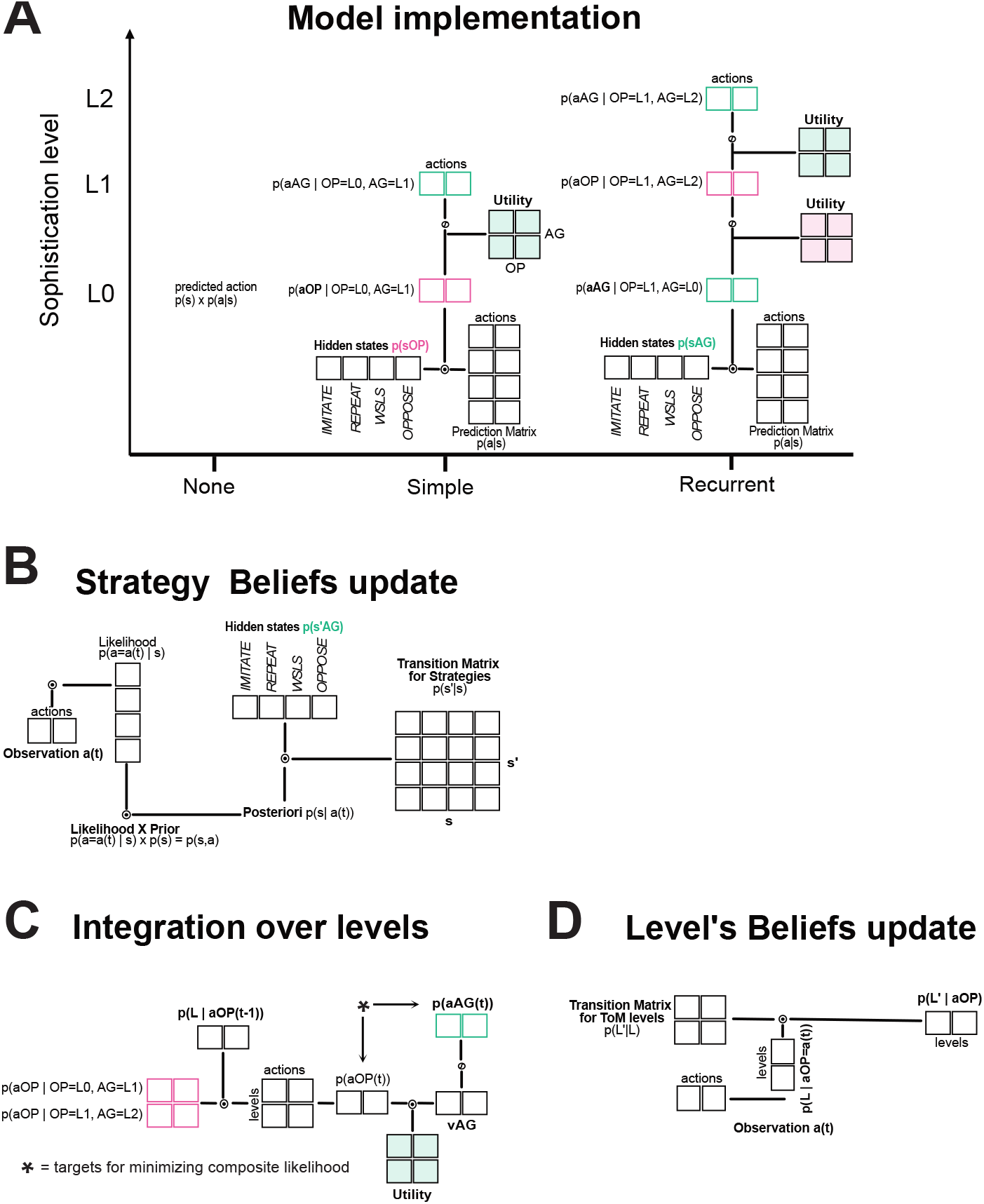
Overview of the BELIEFS model architecture. (A) Model implementation. At the base level, the model maintains a belief distribution over the Opponent’s Level-0 strategies. Each strategy deterministically predicts the next action based on the previous observation. These strategy-specific predictions are weighted by their current beliefs to produce the integrated Level-0 action prediction for the Opponent. One level higher, the model generates a Level-1 Agent that best responds to these Level-0 predictions using its utility matrix. A Level-2 Agent is formed recursively, reasoning about an Opponent who is itself using a Level-1 best response. **(B) Updating Level-0 strategy beliefs**. Beliefs about the Opponent’s L0 strategies are updated using the same Bayesian HMM mechanism. The model evaluates how well each strategy’s predicted action accounts for the observed choice, updates the strategy posterior accordingly, and then applies a strategy-transition matrix that governs the probability of persisting in or switching between strategies. The resulting distribution serves as the prior for the next trial. **(C) Integration over ToM levels (BELIEFS)**. BELIEFS combines the Opponent’s action predictions across ToM levels (e.g., L0 and L1) by weighting each level’s predictions with the current belief over the Opponent’s ToM level. These weighted predictions are normalized and collapsed across levels, yielding a single integrated prediction of the Opponent’s next action. This integrated prediction is used by a higher-level Agent (e.g., L2) to compute its best response. **(D) Updating beliefs about ToM level**. Beliefs about the Opponent’s ToM level are updated via a second HMM process. After observing the Opponent’s action, the model evaluates the likelihood of that action under each ToM level, updates the level beliefs, and applies a ToM-level transition matrix that specifies the probability of remaining at or switching between ToM levels. This transition-adjusted belief distribution becomes the prior for the next trial.

## Acknowledgment

We thank Oleg Solopchuk for his valuable suggestions, guidance on the model development, and insightful contributions during internal discussions. This work was funded by grants the German Research Council (DFG) to J.G., namely the Collaborative Research Center SFB 1528 “Cognition of Interaction” and the Research Unit RU5389 “Dynamic Belief Updating”.

## Credit authorship contribution statement

G.L.M.; Conceptualization, Writing - original draft, Software, Analysis, Visualization. S.K.; Conceptualization. J G; Conceptualization, Funding acquisition, Writing - original draft, Software.

## Footnotes

1 Note that *p*(*a*_*OP*_(*t*)) refers to the action set of the agent at trial *t* (i.e. a belief distribution over all available action), while *c*_*OP*_(*t*) refers to a specific choice in trial *t*.

